# RNA-based sensitive fungal pathogen detection

**DOI:** 10.1101/2023.09.26.559494

**Authors:** Julia Micheel, Franziska Aron, Abdulrahman A. Kelani, Christian Girbardt, Matthew G. Blango, Grit Walther, Damian Wollny

## Abstract

Detecting fungal pathogens, a major cause of severe systemic infections, remains challenging due to the difficulty and time-consuming nature of diagnostic methods. This delay in identification hinders targeted treatment decisions and may lead to unnecessary use of broad-spectrum antibiotics. To expedite treatment initiation, one promising approach is to directly detect pathogen nucleic acids such as DNA, which is often preferred to RNA because of its inherent stability. However, a higher number of RNA molecules per cell makes RNA a more promising diagnostic target which is particularly prominent for highly expressed genes such as rRNA. Here, we investigated the utility of a minimal input-specialized reverse transcription protocol to increase diagnostic sensitivity. This proof-of-concept study demonstrates that fungal rRNA detection by the minimal input protocol is drastically more sensitive compared to detection of genomic DNA even with high levels of human RNA background. This approach can detect several of the most relevant human pathogenic fungal genera, such as *Aspergillus, Candida*, and *Fusarium* and thus represents a powerful, cheap, and easily adaptable addition to currently available diagnostic assays.

## Introduction

Invasive fungal infections are an emerging health problem causing millions of life-threatening conditions worldwide, as highlighted in the recent release of a World Health Organization (WHO) Fungal Priority Pathogens List.^1^ Although often overlooked, the number of fatal fungal infections is estimated to be more than 1.5 million per year^2^, with the main reason for a continued rise in global fungal infections attributed to an increase in susceptible, immunocompromised patients. Particularly vulnerable are solid-organ transplant and hematopoietic stem-cell recipients or patients reliant on other immune-modifying drugs.^3,4^ Four fungal genera cause the majority of these deadly invasive infections, *Aspergillus, Candida, Cryptococcus*, and *Pneumocystis*, typically via environmental exposure, but shifts and expansions in the prevalence of endemic fungi have also been observed and attributed to human migration, climate change, and even the use of antifungals in agriculture, among others.^5^ This makes pathogen identification in areas of non-endemicity challenging due to a lack of clinical expertise.^6,7^

Early detection of fungal infections is crucial for rapid treatment initiation and consequently successful therapy.^4^ However, at present therapy decisions are made on pathogen identification frequently based on cultivation, which is especially time consuming for fungal cultures.^8^ Cultivation of fungal pathogens takes several days to weeks and is not always successful due to challenging culture conditions or contamination with faster growing microorganisms such as (commensal) bacteria.^8,9^ Especially *Candida* and *Aspergillus* blood culture sensitivity is low with success rates only up to 50%.^8–10^ Alternative methods such as histopathology and serological tests for particular species such as the galactomannan test restricted to aspergillosis are faster but lack sensitivity. Treatment decisions are then delayed and must often be performed based on the clinical examination only.^8^ A further complication is that, in many cases, discrimination between bacterial and fungal infection is not possible based on symptoms alone.^8,9^

The detection of fungal nucleic acids directly from biopsies represents a promising route to circumvent the tedious and unreliable cultivation process. Compared to pathogen cultivation, direct DNA isolation and subsequent PCR is considerably faster, producing results in hours instead of days or weeks.^8,9,11–13^ In comparison to DNA, RNA molecules are present at higher copy numbers in cells. Thus, targeting RNA molecules for diagnosis should increase the sensitivity of the assay.^14^ This is particularly true for highly expressed genes such as e.g. ribosomal RNA (rRNA).^15^

Although working with RNA is traditionally considered challenging due to its fragile nature, technical advancements in the last decade enhanced the promise of RNA-based diagnostics. In particular the field of single-cell RNA sequencing has demonstrated that a lot of valuable information can be obtained even from minimal amounts of RNA.^16,17^ Specifically, the protocols developed in this field advanced reverse transcription efficiency e.g. by utilization of optimized enzymes, adapted buffer conditions, addition of other reagents like crowding reagents, longer incubation times, and thermal cycling to unfold RNA secondary structures.^18,19^ In addition, a template-switching mechanism for cDNA second strand synthesis was implemented to ensure 5’ coverage and enables subsequent cDNA amplification.^20^ Furthermore, the reaction volumes are low, which leads to a high concentration of the output cDNA for further applications and significantly decreases the cost per reaction.^19,21^

Using highly abundant target RNA, like rRNA, with such optimized methods should enable fast and sensitive detection of fungal RNA which, in the future, could be utilized for rapid diagnosis and subsequent therapy initiation. Here, we demonstrate RNA-based fungal pathogen detection by application of a specialized minimal input reverse transcription protocol adapted from SMART-seq3^19^ for efficient cDNA synthesis and subsequent qPCR. We find that our method is more sensitive to detection of fungal-derived nucleic acids compared to standard DNA and RNA (RT-)qPCR protocols. This proof-of-concept study demonstrates that technical advances from the single-cell transcriptomics field might help to increase detection sensitivity for diagnostics and beyond.

## Material and methods

### Input materials

#### Fungal culture conditions

*Aspergillus fumigatus* (*A. fumigatus*) wild type (CEA17Δ*akuB*^*KU80*^) were grown on *Aspergillus* minimal medium (AMM) agar (1.5%) plates at 37°C for 5 days (70 mM NaNO_3_, 11.2 mM KH_2_PO_4_, 7 mM KCl, 2 mM MgSO_4_, 1% (w/v) glucose and 1 μl/ml trace element solution (pH 6.5)). The trace element solution was composed of 1 g FeSO_4_ • 7 H_2_O, 8.8 g ZnSO_4_ • 7 H_2_O, 0.4 g CuSO_4_ • 5 H_2_O, 0.15 g MnSO_4_ • H_2_O, 0.1 g NaB_4_O_7_ • 10 H_2_O, 0.05 g (NH_4_)_6_Mo_7_O_24_ • 4 H_2_O, and ultra-filtrated water to 1000 ml. Mycelium were grown in liquid AMM culture for 24 h at 37°C shaking at 200 rpm with an initial inoculum of 10^8^ fungal conidia.

For genomic DNA isolation, roughly 0.4 g of ground mycelial biomass was transferred into 2-ml microcentrifuge tubes and suspended in 450 μl of Cell Lysis Solution (1M Tris (pH 7), 10% SDS, and 0.5 M EDTA (pH 8)) before incubation at 65°C for 15 min. Samples were placed on ice for 5 min, 300 μL of 5 M ammonium acetate was added and samples were vortexed for 10 s. Cellular debris was collected by centrifugation for 10 min at ≥10,000 rpm and the supernatant transferred to clean microcentrifuge tubes. DNA was precipitated by mixing supernatant with 500 μl of isopropanol and incubating at room temperature for at 20 min. DNA pellets were then collected by centrifugation for 10 min at ≥10,000 rpm. Supernatant was discarded and DNA pellets washed with 500 μl of 70% ethanol by centrifugation for 3 min at ≥10,000 rpm. Ethanol was removed by pipetting and discarded. DNA pellets were allowed to air dry for 5-10 min before being resuspended in 50-100 μL of MilliQ water. DNA samples were stored at -20°C.

*A. fumigatus* RNA was isolated as described previously.^22^ Briefly, fungal mycelium grown in liquid culture was collected using Miracloth (Millipore, USA) and disrupted in liquid nitrogen using a precooled mortar and pestle. Approximately 0.5 g of homogenized fungal mycelium was transferred into a 2-ml Eppendorf tube. 800 μl TRIzol was then added to the disrupted mycelia and vortexed vigorously. Tubes were frozen briefly in liquid nitrogen for 5 sec and allowed to thaw, while kept on ice. 160 μl of chloroform was added, samples were vortexed, and then centrifuged for 5 min at 4°C at full speed. Without disturbing the interphase, the aqueous upper phase was transferred to a fresh 2-ml tube. RNA extraction from aqueous phase was done with 1 volume of phenol/chloroform/isoamyl alcohol (25:24:1, v/v/v). After brief vortexing, samples were centrifuged for 5 min at 4°C. The extraction was repeated until no more interphase was observable, followed by another extraction with 400 μl chloroform. RNA was precipitated using 400 μl isopropanol for 20 min, followed by centrifugation for 20 min at 4°C. The pellet was washed with 700 μl 70% ethanol, air-dried at 37°C for 5 min, and resuspended in RNase-free water. DNase treatment was then performed on the samples using 2 units of TURBO DNase (Thermo Fisher Scientific) per 10 μg RNA for 30 min at 37°C in 100 μl total volume. Finally, Total RNA was collected using the RNA Clean and Concentrator-25 kit (Zymo Research, USA) according to the manufacturer’s instructions.

For DNA extraction of *Candida* and *Fusarium* species, filamentous fungi were grown on 4% malt extract agar (BD Difco, USA), yeasts were grown on yeast extract peptone dextrose agar (1% yeast extract (Serva, Germany), 2% peptone (BD Difco), 2% glucose (Carl Roth)). Genomic DNA was extracted from 2-to 5-day-old cultures by two different protocols. For both protocols, fungal material was transferred to a tube containing acid-washed glass beads and 1 ml of lysis buffer (50 mM Tris, 50 mM sodium EDTA, 3% (w/v) sodium dodecyl sulfate (SDS) (pH 8)). The samples were homogenized for 5 min at maximum speed using a vortex adapter, followed by 1 h of incubation in a thermomixer at 68°C. Thereafter, the tubes were spun for 10 min at 16,000 relative centrifugal force (RCF), and the supernatant was transferred to a new 2-ml tube. Applying the first protocol, an equal volume of a mixture of phenol, chloroform, and isoamyl alcohol (25:24:1, v/v/v (pH 7.5 to 8.0)) was added. The samples were mixed by turning and spun for 10 min at 16,000 RCF. The upper (aqueous) phase was transferred to a new tube, and the step was repeated. Then, a 0.5 volume of 99.9% ethanol was added to precipitate the DNA. Using the second protocol 225 μl of 3 M sodium acetate were added to the supernatant and mixed by multiple inversions. After incubation of 10 min at -20°C the samples were spun for 10 min at 16,000 RCF and the supernatant was transferred to a new 2 ml tube. Then 1 ml EtOH was added, mixed by multiple inversions and incubated for at least 20 min at -20°C to precipitate the DNA. In both protocols the DNA was pelleted at 16,000 RCF for 10 min. The supernatant was decanted, and the DNA pellet was washed twice with 200 ml of 70% ethanol, dried, resuspended in 50 μl of distilled water, and stored at -20°C.

#### Human cell culture conditions

RPE-1 hTERT cells (ATCC) were cultured in DMEM:F12 media (Thermo Fisher Scientific, Germany) and culture media were supplemented with 10% fetal bovine serum (FBS; Thermo Fisher Scientific) and 1% penicillin/streptomycin (Thermo Fisher Scientific). For nucleic acid isolation, cells were washed with PBS (Gibco, USA) and trypsinized (Gibco). DNA isolation was performed using the DNeasy Blood and Tissue kit (Qiagen, Germany) and for RNA extraction, the RNeasy Plus Mini kit (Qiagen) was used, both according to the manufacturer’s protocols.

#### Bacterial culture conditions

*Escherichia coli (E. coli)* strain B (Migula) Castellani and Chalmers (ATCC, USA) was cultivated in LB medium (Luria/Miller) (Carl Roth, Germany) at 37°C and 200 rpm overnight. RNA isolation was performed using TRIzol Reagent (Invitrogen, USA) as described in the manufacturer’s protocol.

For DNA isolation, an in-house protocol for proteinase K treatment and subsequent phenolchloroform-isoamyl alcohol DNA extraction was performed. In detail, an *E. coli* cell pellet from a 2 ml culture was resuspended in 580 μl proteinase K buffer (containing 100 μl PBS, 0.005% SDS (Carl Roth), and 60 μg proteinase K (Roche, Switzerland) in TE buffer (PanReac AppliChem, Germany)). The mixture was incubated at 50°C and 400 rpm for 1 h. Afterwards, 80 μl CTAB/NaCl solution (10% CTAB in 0.7 M NaCl (both Carl Roth)) was added, mixed, and incubated at 65°C for 15 min. One volume of phenol-chloroform-isoamyl alcohol mix (25:24:1, v/v/v) was added, the mixture was inverted, and centrifuged for 5 min at full speed. The supernatant was transferred into a fresh tube. Again, one volume of phenol-chloroform-isoamyl alcohol mix (25:24:1, v/v/v) was added, the mixture inverted, and centrifuged for 5 min at full speed. For DNA precipitation, sodium acetate (Sigma Aldrich) solution was added to the supernatant to a final molarity of 0.3 M. 4 volumes of 100% EtOH were added and the mixture was inverted. The DNA solution was incubated for 2 h at -20°C and afterwards centrifuged at full speed for 30 min at 4°C. The pellet was washed twice with 80% EtOH, air dried and dissolved in nuclease-free water.

### SMART-seq3-adapted reverse transcription

The protocol for reverse transcription has been adapted from SMART-seq3^19^ while oligonucleotides were adapted from SMART-seq2^18^. The concentration of the total RNA input was measured using the Qubit 3 Fluorometer (Thermo Fisher Scientific) with the Qubit RNA High Sensitivity assay kit (Invitrogen) and diluted with nuclease-free water to the specified concentration. The input RNA (in 1 μl nuclease-free water) was added to 3 μl lysis mix (containing 5% PEG 8000 (Carl Roth), 0.1% Triton X-100 (Roche), 1.6 U Recombinant RNase Inhibitor (Takara, Japan), 0.5 μM random hexamer (IDT, USA, sequence shown in Table 1), and 0.5 mM dNTPs (Thermo Fisher Scientific). The lysis mix was incubated for 10 min at 72°C. Afterwards, 1 μl RT mix was added (containing 25 mM Trizma hydrochloride solution pH 8 (Sigma Aldrich), 30 mM sodium chloride (Carl Roth), 2.5 mM magnesium chloride (Invitrogen), 1 mM GTP (Thermo Fisher Scientific), 8 mM DTT (Invitrogen), 0.5 U Recombinant RNase Inhibitor, 2 U Maxima H minus (Thermo Fisher Scientific) and 2 μM template switching oligo (TSO) (IDT, sequence shown in Table 1)). The reverse transcription was performed for 90 min at 42°C, followed by 10 cycles of 2 min at 50°C and 2 min at 42°C each, and an inactivation step of 5 min at 85°C.

**Table 1:**
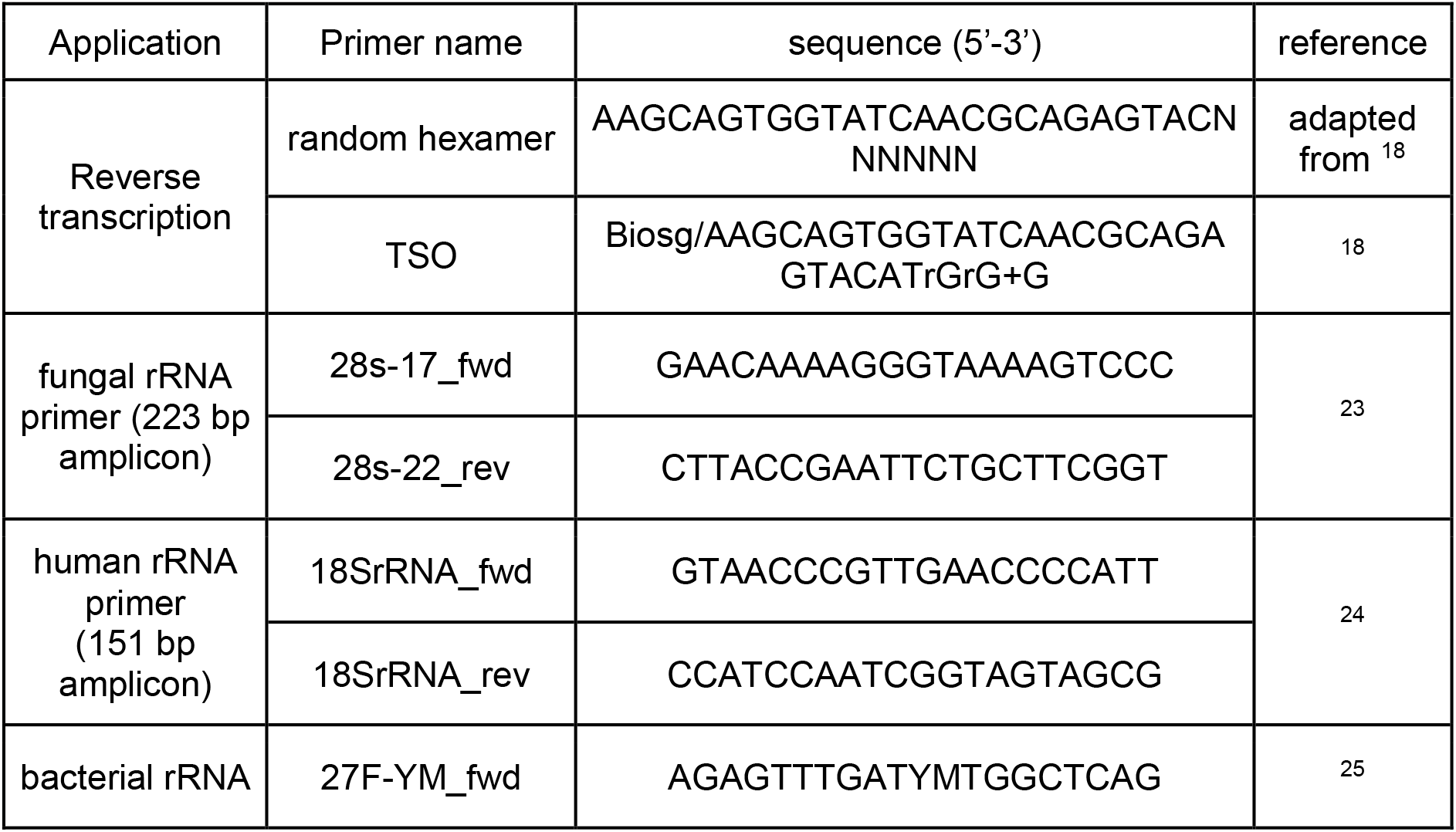
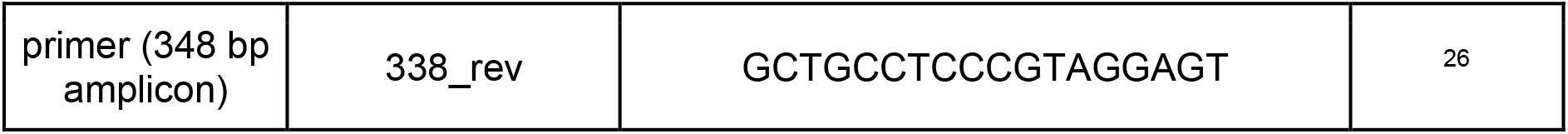
DNA oligos used for reverse transcription and PCR with their respective sequences and original publications. Biosg/ indicates 5’ biotinylation, rG is the RNA-base guanine, +G is a locked guanine base, Y is cytosine or thymine, M is adenine or cytosine.

### Standard reverse transcription

Standard reverse transcription was performed according to the Maxima H minus reverse transcriptase manufacturer’s protocol. The concentration of the total RNA input was measured using the Qubit 3 Fluorometer (Thermo Fisher Scientific) and the Qubit RNA High Sensitivity assay kit (Invitrogen) and diluted with nuclease-free water to the specified concentration. The RNA was added to 14.5 μl master mix (containing 13.8 μM random hexamer (IDT, sequence shown in Table 1) and 0.69 mM dNTPs (Thermo Fisher Scientific). The mixture was incubated at 65°C for 5 min and chilled on ice for 2 min afterwards. 5.5 μl of master mix 2 were added (containing 1x Maxima Buffer (Thermo Fisher Scientific), 20 U Recombinant RNase Inhibitor, and 100 U Maxima H minus reverse transcriptase). Reverse transcription was carried out by 10 min incubation at 25°C, 30 min at 42°C and inactivation for 5 min at 85°C.

### RT-qPCR

RT-qPCR was carried out with 4 μl RT mix derived from standard or SMART reverse transcription or with a specified amount of gDNA. gDNA concentrations were measured using the Qubit 3 Fluorometer and the Qubit dsDNA High Sensitivity assay kit (Invitrogen). Input gDNA or cDNA was added to the qPCR master mix containing 1X Colorless GoTaq® Flexi Buffer (Promega, USA), 4 mM magnesium chloride (Promega), 0.4 mM dNTPs (Thermo Fisher Scientific), 0.2 μM of the respective forward and reverse primer each (IDT, sequences shown in Table 1), 0.5 U GoTaq G2 Flexi Polymerase (Promega), and 0.25x SYBR green (Sigma Aldrich). Reactions were performed on a C1000 Touch Thermal Cycler (Bio-Rad, USA). Initial denaturation was carried out for 3 min at 95°C, followed by 30 cycles of 40 s denaturation at 95°C, 30 s annealing at 60°C, and 30 s elongation at 72°C. Afterwards, a melting curve was generated from 65°C to 95°C with a 0.5°C/min increment. The results were analyzed using the Bio-Rad CFX Manager 3.1 program.

### Agarose gel electrophoresis

Agarose gels (2%) were prepared by melting 2 g of agarose (Biozym LE Agarose, Biozym, Germany) per 100 ml TAE running buffer (Sigma Aldrich) containing 2 drops of ethidium bromide (Carl Roth). 1 kb plus ladder (Invitrogen) and samples each mixed with 1x Loading Dye (New England Biolabs, USA) were loaded onto the gel. Gels were run for 30 to 40 min with 130 V. Images were taken using the Molecular Imager Gel Doc XR+ system (Bio-Rad) and the Image Lab Software (Bio-Rad).

### Phylogenetic analysis

Phylogenetic tree generation was performed with phyloT v2 (https://phylot.biobyte.de) on the basis of NCBI taxonomy on 03/2023. The taxonomy identifiers were used instead of a sequence alignment because for some of the species no sequences were available. The NCBI taxonomy identifiers are listed in Supplementary Table 1. The unrooted tree was visualized using the interactive Tree of Life (iTOL).^27^

### Multiple sequence alignment

The alignment of the fungal rRNA primers and the desired amplicon sequences of a variety of human pathogen fungal species was created using MAFFT version 7 with standard settings^28^ and visualized with Jalview 2.11.2.6.^29^ All fungal 28S rRNA sequences were taken from GenBank with the indicated reference numbers shown in Supplementary Table 1.

## Results

First, we aimed to test the hypothesis that a qPCR assay detecting fungal nucleic acids is more sensitive on the RNA level than on the DNA level (Figure 1a). We hypothesized that this effect should be particularly apparent for rRNA, since rRNA is highly expressed in cells and thus any amount of fungal RNA should offer more priming opportunities compared to the same amount of fungal DNA. In order to detect with the highest sensitivity the RNA molecules transcribed from the rRNA gene, we adapted the SMART-seq3 protocol, which was originally designed for detecting minute RNA amounts from single cells (Figure 1b).^19^ The high sensitivity of the protocol is largely based on the conditions for the reverse transcriptase reactions and includes a template switch to ensure sufficient coverage of the 5’-ends (Figure 1b). We probed the sensitivity of both input materials (DNA vs. RNA) in a dilution series spanning two orders of magnitude (5 ng – 50 pg). To ensure comparability between both approaches, we used the same input amounts of total RNA and DNA (Figure 1c). We found that the detection sensitivity was consistently higher using RNA compared to gDNA irrespective of the input amount (Figure 1c). This result indicates that even if the same amount of RNA and DNA is present, detection of fungal-derived nucleic acids is a lot more sensitive on the RNA level. This is particularly promising since *in vivo* the amount of rRNA and rDNA is not equal. Instead, the cellular rRNA amount vastly exceeds the rDNA amount, which would further increase the chance of detection when rRNA is chosen as the qPCR target.

**Figure 1:**
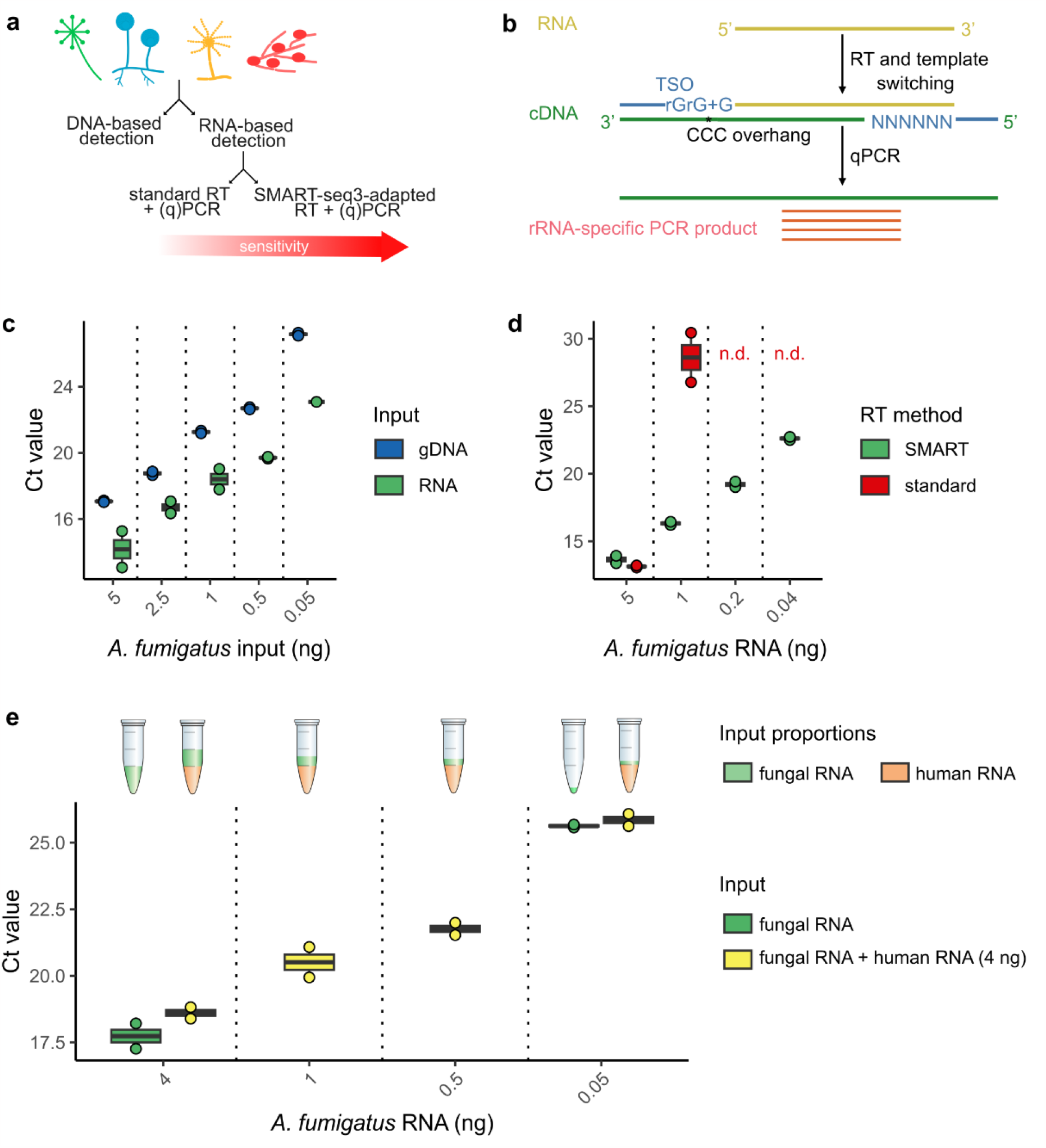
Sensitivity of RNA-based fungal pathogen detection. a. Schematic of the tested quantitative PCR-based pathogen detection methods. RT denotes reverse transcription. **b** Reverse transcription and RT-qPCR workflow. Starting with RNA, SMART-seq3-adapted reverse transcription is carried out including cDNA first-strand synthesis and template switching. Subsequently qPCR is performed for rRNA transcript quantification. The template switching oligo (TSO) including (ribo)guanines(rG/G), and the random hexamer including a 5’-adapter sequence are depicted in blue. The TSO’s rGrG+G bind the complementary CCC overhang at the 3’ end of the newly synthesized cDNA and enable template switching.^19^ **c** Comparison of RT-qPCR detection sensitivity using equivalent amounts of *A. fumigatus* gDNA and *A. fumigatus* total RNA as input material in qPCR (DNA input) or SMART-seq3-based RT and subsequent qPCR (RNA input) in technical duplicates. **d** Comparison of RT-qPCR detection sensitivity using the SMART-seq3-based RT versus a standard RT protocol and equal input amounts of *A. fumigatus* total RNA in technical duplicates. n.d. denotes no detection within 30 PCR cycles. **e** Detection sensitivity of decreasing *A. fumigatus* RNA amounts in the presence of 4 ng human (RPE-1) total RNA in technical duplicates. The RNA ratios within the different samples are visualized in the reaction tubes above.

Next, we tested the detection advantage of the sensitive SMART-seq3 protocol, over a standard reverse transcription protocol for the same RT enzyme (Maxima H-minus reverse transcriptase). RT-qPCR results revealed a clear advantage of the SMART-seq protocol (Figure 1d). When we used the standard reverse transcription according to the manufacturer’s protocol, we were unable to detect input molecules in the picogram range, while the adapted SMART-seq3 protocol reliably detected even small amounts of input RNA (e.g. Ct value of 23 at 40 pg of input RNA). The main reason for this effect is the considerably lower reaction volume for which the SMART-seq3 protocol was optimized. Yet, other factors such as optimization of the salt conditions of the reverse transcription buffer or the addition of molecular crowding reagents likely have also contributed to the high detection efficiency.^19^

When RNA is isolated from patient samples for a diagnostic test, large amounts of human RNA will likely accompany the pathogen RNA that the assay aims to detect. In order to design a reliable test, it is thus crucial that a) the sensitivity is not negatively affected by excessive amounts of human “background” RNA and that b) the primers are highly specific for the pathogen RNA to avoid false positive results. Therefore, we tested the detection sensitivity of fungal RNA by mixing decreasing amounts of fungal RNA (from 4 ng to 50 pg) with a constant, large amount of human RNA (4 ng). We found that the detection sensitivity is not affected by the presence of human RNA, even if almost 100 times more human RNA (4 ng human RNA vs. 50 pg fungal RNA) is present in the reaction (Figure 1e).

Next, we tested the specificity of the fungal detection assay. Here, the primers used for fungal rRNA detection must satisfy two major requirements. On the one hand, the primers must be located in a conserved region of a highly expressed gene in order to detect as many fungal pathogen species as possible. On the other hand, they must not produce false-positive results due to the amplification of human RNA or amplification of other potential pathogenic sources such as e.g. bacterial RNA. Thus, we chose primers that anneal to a coding region of the 28S rRNA, which is conserved among fungal species (Figure 2a). Using human and bacterial RNA for comparison, we found that the primers only amplify fungal rRNA (Figure 2b, Supplementary Figure 1a).

**Figure 2:**
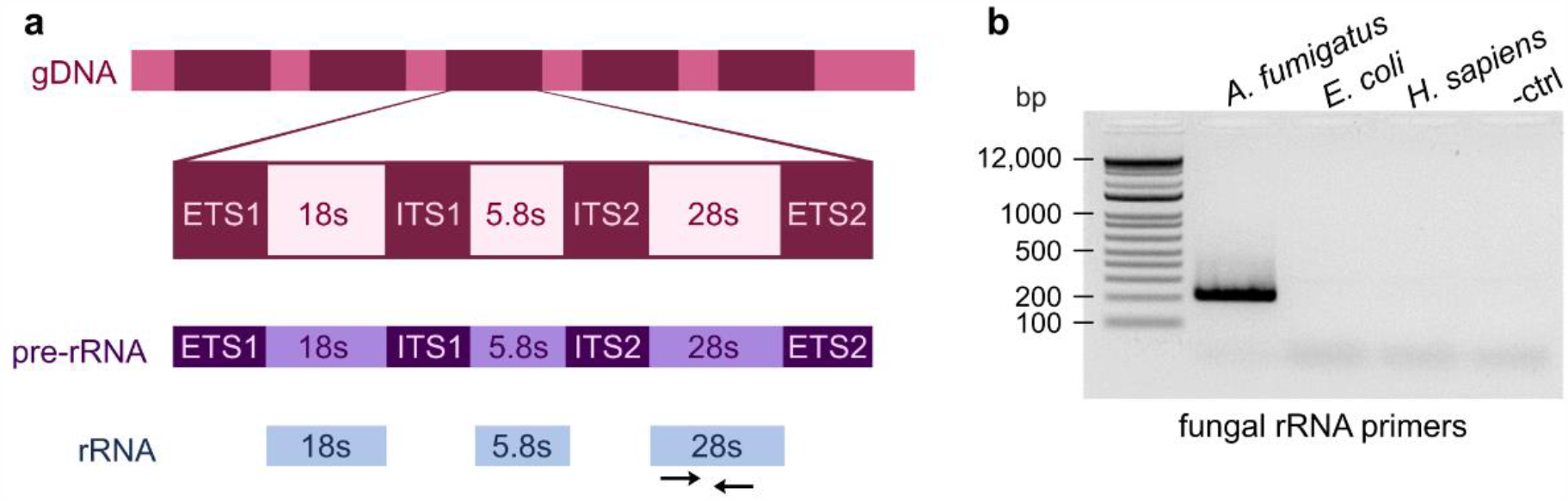
Fungal pathogen detection specificity. **a** Schematic of the primer binding sites on DNA, pre-rRNA, and rRNA level in the fungal genome and transcripts. **b** Primer specificity for fungal rRNA gene detection from *A. fumigatus* DNA versus bacterial (*E. coli* DNA) and human (RPE-1 DNA) input material.

In a clinical setting, a negative result for a fungal rRNA detection assay may have several reasons. A trivial explanation would be that the PCR assay failed. Since human RNA will likely be present at high amounts in human samples, we set up human-specific rRNA primers that are not cross-reactive with fungal or bacterial RNA as a positive control for the PCR (Supplementary Figure 1b, Supplementary Figure 2). Fungal RNA produced a faint band of the desired size in the agarose gel (Supplementary Figure 2) and a weak signal in the qPCR (Supplementary Figure 1b) below the detection threshold. Another reason is that the infection is caused by a different pathogen such as bacteria instead. Thus, we aimed to broaden the scope of our detection assay beyond the fungal kingdom.

To detect bacterial pathogens, we used the universal bacterial rRNA primers 27F-YM_fwd^25^ and 338_rev^26^. As with the detection of fungal rRNA, bacterial RNA consistently exhibited greater sensitivity than the same quantity of bacterial DNA (Supplementary Figure 3a). This effect was particularly pronounced at low RNA/DNA amounts (Supplementary Figure 3a). The primers are specific for bacterial input material as no false-positive PCR products were detected with human or fungal input material (Supplementary Figure 3b). Again, human and fungal RNA produced a faint band of the desired size in the agarose gel (Supplementary Figure 3b) and a weak signal in the qPCR (Supplementary Figure 1c) below the detection threshold highlighting the need of a Ct cut off to avoid false-positive results. Similar to the detection of fungal RNA, the detection sensitivity was not negatively affected even by the presence of high amounts of human RNA (Supplementary Figure 3c).

**Figure 3:**
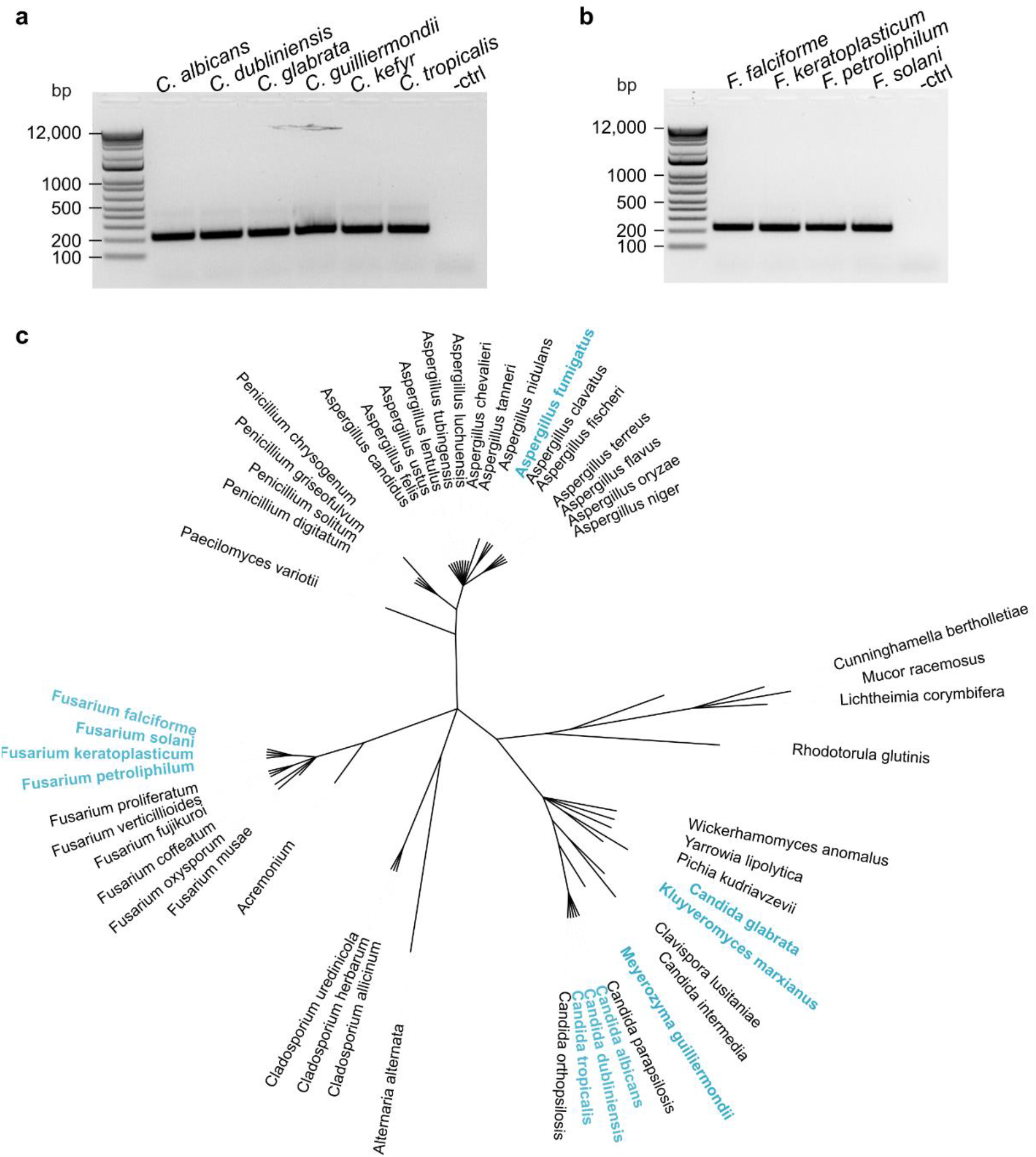
Fungal PCR primer conservation. PCR products for different **a** *Candida* or **b** *Fusarium* species using the fungal rRNA primers. DNA was used as input material for PCR analysis. **c** Phylogenetic tree depicting the fungal species that can be detected using the fungal rRNA primers. Computational and/or experimental confirmations are depicted in black or blue respectively. The phylogenetic tree was generated using phyloT (03/2023) based on the NCBI taxonomy identifiers (Supplementary Table 1) and visualized using iTol.

A third reason for a negative assay result would be that the primer sequence is not conserved across all potential fungal pathogens. Based on *in silico* analysis, we found that the primers we used are intended to detect the most prevalent pathogenic species known to cause severe fungal infections which is consistent with previous results.^23^ In order to validate the computational analysis experimentally, we tested our fungal detection assay beyond *A. fumigatus* on six *Candida* (Figure 3a) and four *Fusarium* species (Figure 3b). In agreement with the computational conservation analysis (Figure 3c), all tested fungal species could be detected with our assay indicating that the assay is reliable across a wide phylogenetic span.

## Discussion

Given the increasing prevalence of fungal infections^3,4,30^, there is a growing need for rapid and reliable pathogen detection methods in order to initiate targeted therapy as early as possible. This is particularly true for diseases where the pathogen prevalence in the body is low and thus technically challenging and time-consuming to detect. Depending on the infection site and the stage of infection, sometimes only small amounts of the pathogen are found. For example, during candidemia, a *Candida* bloodstream infection, as few as 1 colony forming unit was detected per ml blood.^31^ Therefore, it is crucial that detection methods have a high sensitivity. Considering the differences between the amount of RNA and DNA in a cell, the advantage of RNA detection over DNA can be decisive for successful diagnosis.^14^ Conversely, for specimens where more pathogens are present, more sensitive protocols can reduce the amount of biopsy that needs to be taken. Especially in diseases such as eye infections, where biopsy is painful and possibly debilitating, patients can benefit greatly from sampling reduction.

The stability of RNA and consequently its degradation might be a concern resulting in higher reluctance to rely on RNA-based detection assays in the clinical setting. Unlike DNA, RNA requires storage at -80°C, urgent avoidance of freeze-thaw cycles, and RNase-free processing. In addition, the protocols are more complex and longer due to the additional step of cDNA synthesis. However, in our opinion, these disadvantages are outweighed by the diagnostic advantage. Protocols for handling RNA are improved constantly, in particular by the fields of single-cell transcriptomics and liquid biopsy.^21,32,33^ Thus, diagnostic applications can also benefit from optimizations, particularly by the single-cell field. The implementation of these methods is cheap in regard to the low amounts of expensive reagents used (e.g. recombinant proteins) and customized buffers which could foster the implementation of the technique in routine clinical labs.

The primers currently used for our assay are able to detect many human pathogen fungal species of the genera *Aspergillus, Candida, Fusarium*, and *Penicillium*, which are prominent sources of invasive human infections, causing e.g. lung infection^34^, sepsis^35,36^ or eye-related infections.^37,38^ These genera are representatives of both yeasts and molds, which shows the high conservation of the genomic regions that are targeted by the primers (Figure 3, Supplementary Figure 4). Thus, an assay with these primers can enable a rapid initial differentiation between bacterial and fungal infection in order to start a targeted therapy. In the future, with advancements in primer design, a differentiation of different genera could also be performed in order to adjust therapy decisions. Furthermore, we have shown that the assay is suitable for sensitive detection of bacterial rRNA demonstrating potential application of the assay beyond fungal pathogen detection. The increased sensitivity of RNA not only applies to rRNA detection, but can also be used for the detection of protein-coding genes. This is an exciting possibility, especially for the detection of resistance genes or resistance-causing mutations. For instance, in *Candida glabrata* an acquired mutation in the FKS1 and FKS2 genes leads to resistance to echinocandins which are first-line treatment for *Candida* infections.^39^ On the DNA level only two copies of these genes are encoded per cell, while many transcripts are expressed. Similar phenotypes for azole resistance have been reported in *A. fumigatus*, where RNA-based methods could be used to detect Cyp51A overexpression indicating drug resistance.^40^

A limitation of molecular pathogen detection is that with increasing sensitivity there is always a risk of contamination, especially with amplification-based methods. Therefore, negative controls and Ct-cut offs are inevitable to avoid false-positive results. Furthermore, distinguishing between commensal colonization and active infection is difficult with the detection of rRNA genes or protein-coding housekeeping genes.^8^ If genes whose expression marks the transition from commensal to active infection could be identified for future studies, they could be detected on an RNA basis. This would be an additional advantage (besides sensitivity) of using RNA over DNA because, in contrast, at the DNA level, all genes in the genome are detected regardless of whether they are expressed.

In summary, we have demonstrated the advantages of RNA-based approaches for the detection of fungal pathogens. This approach exhibits high sensitivity, is easy to implement, and is expandable for additional species or target RNAs. Therefore, it has the potential to serve as a valuable addition to the existing toolkit for detecting fungal pathogens.

## Data availability

The data supporting the findings of this study are available from the corresponding authors upon reasonable request.

## Author Contributions Statement

J.M., C.G. and D.W. conceptualized the experiments; J.M. and F.A. performed the experiments; A.A.K., M.G.B. and G.W. provided experimental materials; J.M., D.W., M.G.B., A.A.K. and G.W. wrote the manuscript; all authors reviewed the manuscript.

## Acknowledgements

J.M., F.A. and D.W. were supported by the Ministry for Economics, Sciences and Digital Society of Thuringia under the framework of the Landesprogramm ProDigital (DigLeben-5575/10-9) and thurAI (2021 FGI 0009). A.A.K. and M.G.B. were supported by the Federal Ministry for Education and Research (BMBF: https://www.bmbf.de/), Germany, Project FKZ 01K12012 “RFIN – RNA-Biologie von Pilzinfektionen”. The funders had no role in the study design, data collection and analysis, decision to publish, or preparation of the manuscript.

