## Supplementary Material for "RNA-based sensitive fungal pathogen detection"

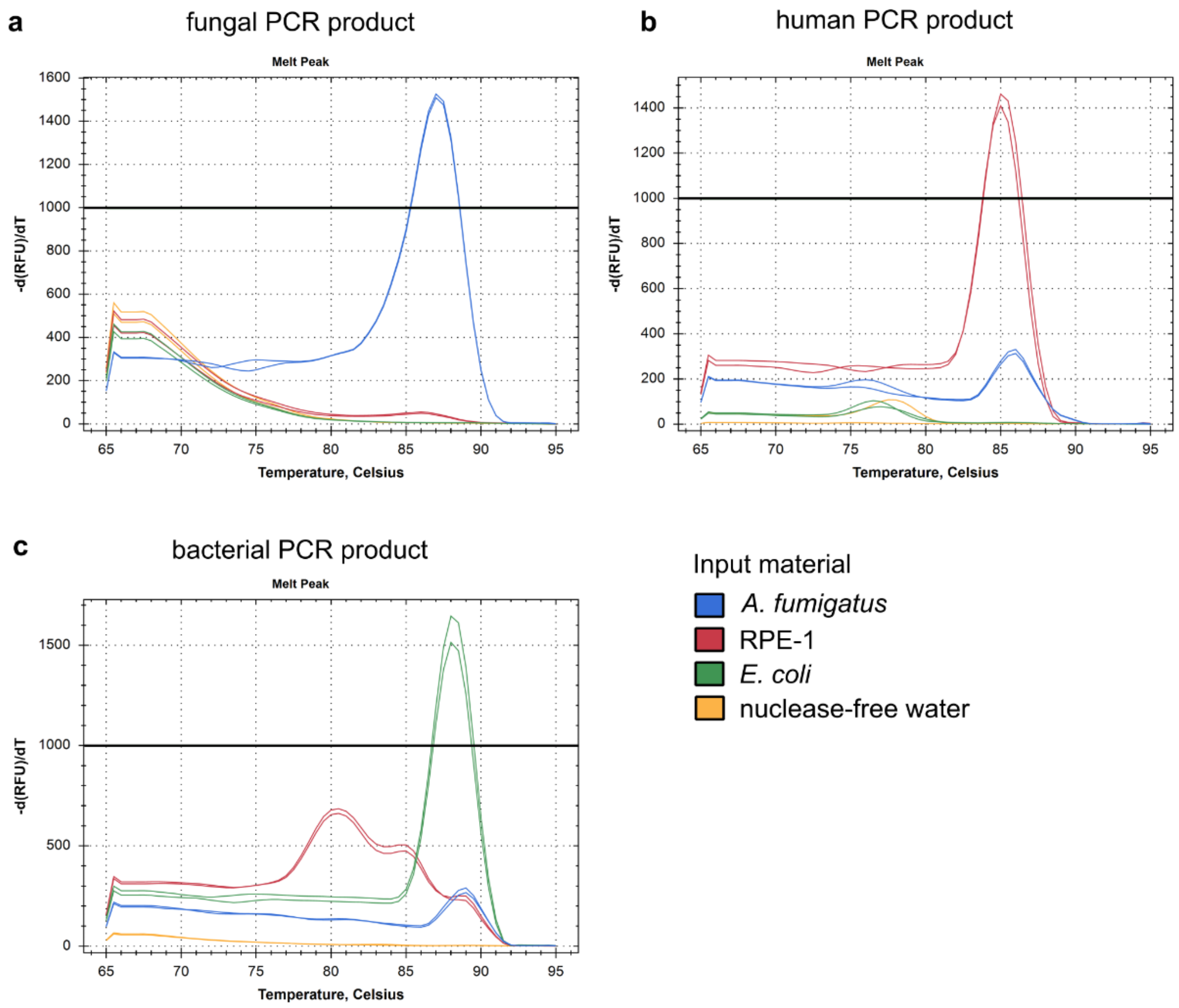


Supplementary Figure 1: **qPCR product melt peaks.** 5 ng gDNA of each indicated species were used as input material for testing the **a** fungal, **b** human and **c** bacterial rRNA primers in technical duplicates. After 30 PCR cycles, melt peaks of the products were generated between 65°C and 90°C with a 0.5°C/min increment.

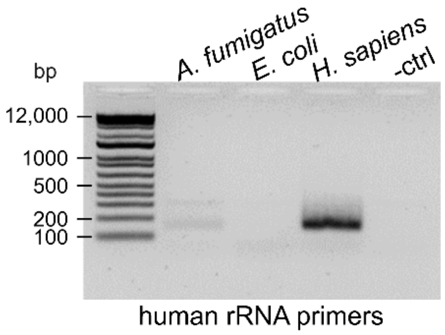


Supplementary Figure 2: **Human PCR primer specificity.** Primer specificity for human rRNA gene detection from RPE-1 DNA versus bacterial (*E. coli* DNA) and fungal (*A. fumigatus* DNA) input material. 5 ng each were analyzed by RT-qPCR and subsequent agarose gel electrophoresis.

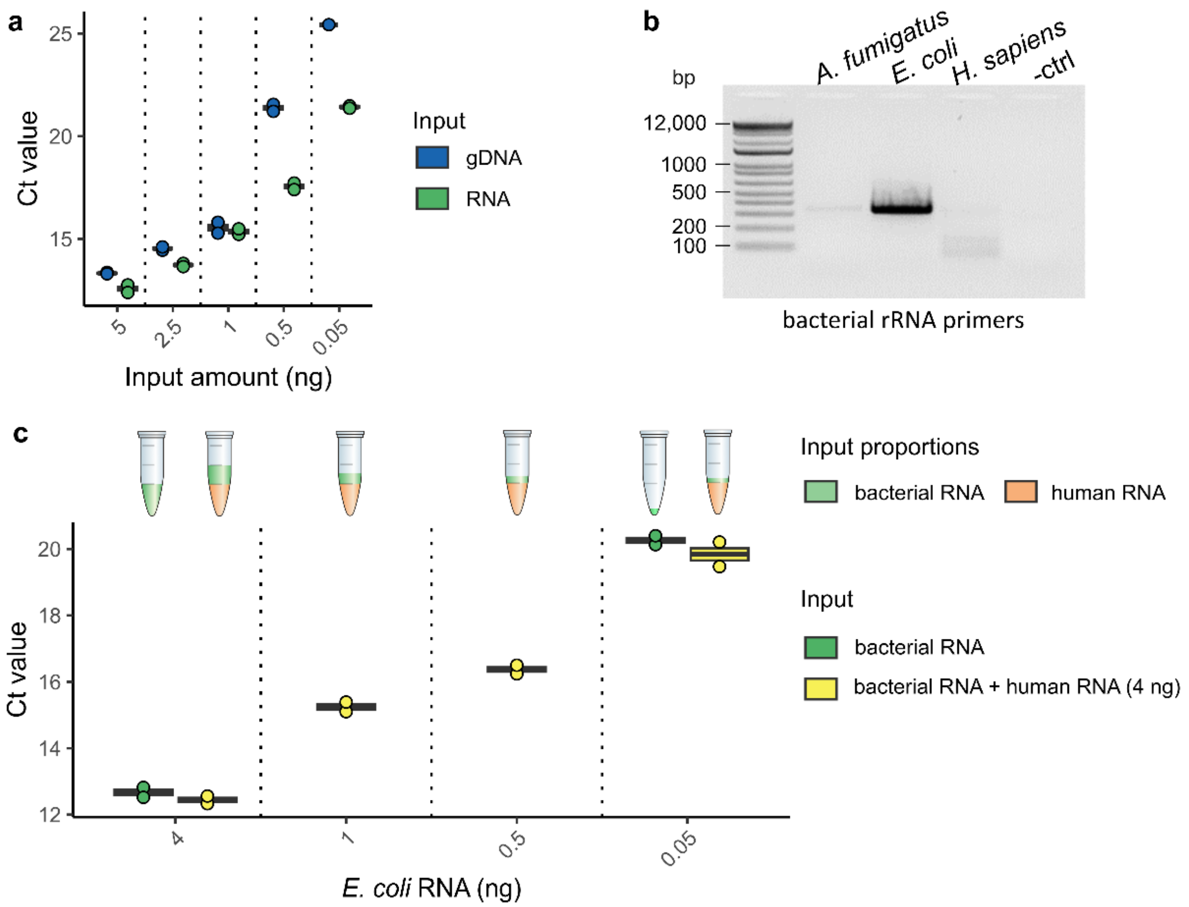


Supplementary Figure 3: **RNA-based bacterial pathogen detection. a** Comparison of RT-qPCR detection sensitivity using equal amounts of *E. coli* gDNA and *E. coli* total RNA as input material in qPCR (DNA input) or SMART-seq3-based RT and subsequent RT-qPCR (RNA input) in technical duplicates. **b** Primer specificity for bacterial rRNA gene detection from *E. coli* DNA versus fungal (*A. fumigatus* DNA) and human (RPE-1 DNA) input material. **c** Detection sensitivity of decreasing *E. coli* RNA amounts in the presence of 4 ng RPE-1 total RNA in technical duplicates. The RNA ratios within the different samples are visualized in the reaction tubes above.

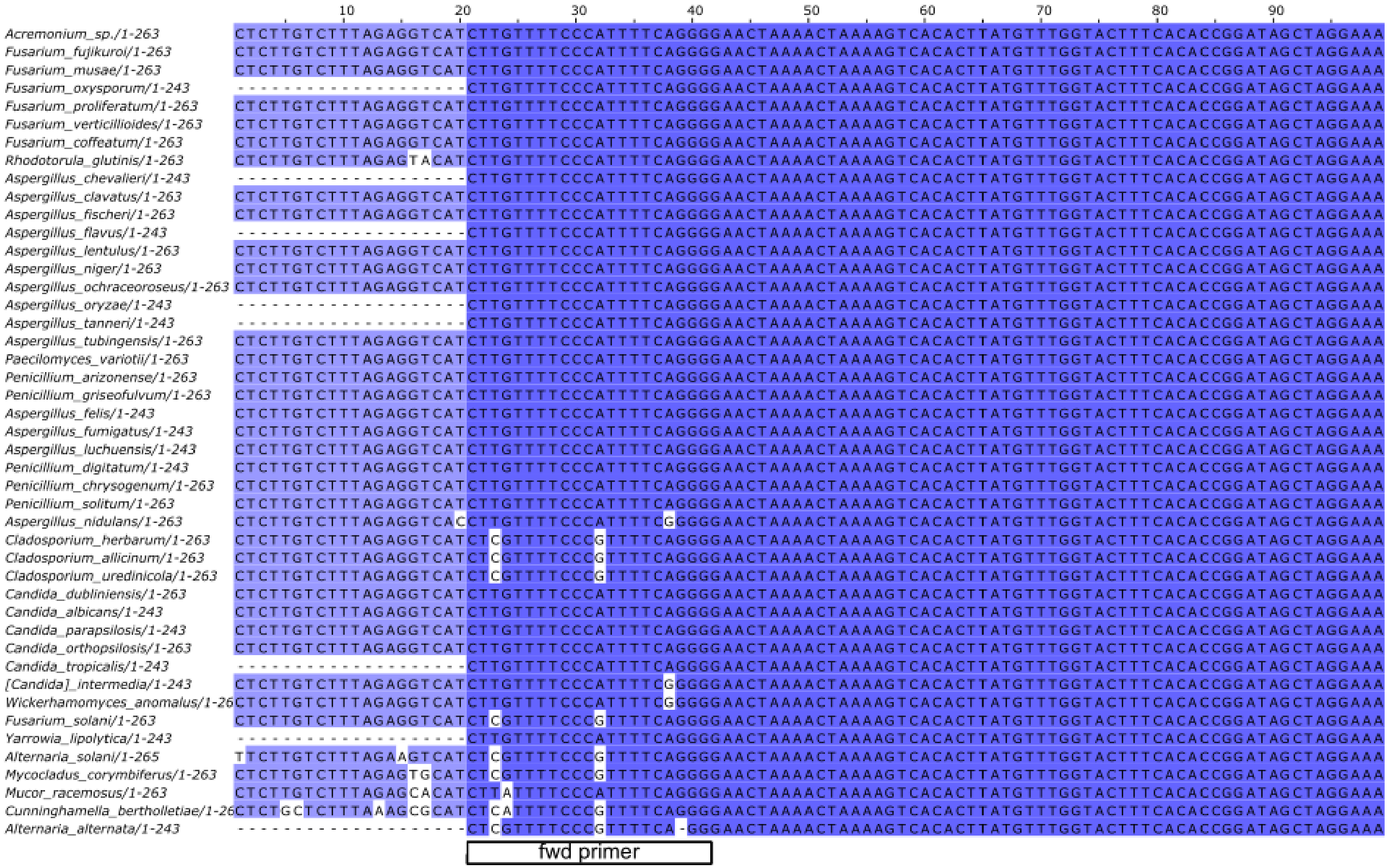
Acremonium_sp./1-263
Fusarium_fujikuroi/1-263
Fusarium_musae/1-263
Fusarium_oxysporum/1-243
Fusarium_proliferatum/1-263
Fusarium_verticillioides/1-263
Fusarium_coffeatum/1-263
Rhodotorula_glutinis/1-263
Aspergillus_chevalieri/1-243
Aspergillus_clavatus/1-263
Aspergillus_fischeri/1-263
Aspergillus_flavus/1-243
Aspergillus_lentulus/1-263
Aspergillus_niger/1-263
Aspergillus_ochraceoroseus/1-263
Aspergillus_oryzae/1-243
Aspergillus_tanneri/1-243
Aspergillus_tubingensis/1-263
Paecilomyces_variotii/1-263
Penicillium_arizonense/1-263
Penicillium_griseofulvum/1-263
Aspergillus_felis/1-243
Aspergillus_fumigatus/1-243
Aspergillus_luchuensis/1-243
Penicillium_digitatum/1-243
Penicillium_chrysogenum/1-263
Penicillium_solitum/1-263
Aspergillus_nidulans/1-263
Cladosporium_herbarum/1-263
Cladosporium_allicinum/1-263
Cladosporium_uredinicola/1-263
Candida_dubliniensis/1-263
Candida_albicans/1-243
Candida_parapsilosis/1-243
Candida_orthopsilosis/1-263
Candida_tropicalis/1-243
[Candida]_intermedia/1-243
Wickerhamomyces_anomalus/1-26
Fusarium_solani/1-263
Yarrowia_lipolytica/1-243
Alternaria_solani/1-265
Mycocladus_corymbiferus/1-263
Mucor_racemosus/1-263
Cunninghamella_bertholletiae/1-26
Alternaria_alternata/1-243
fwd primer

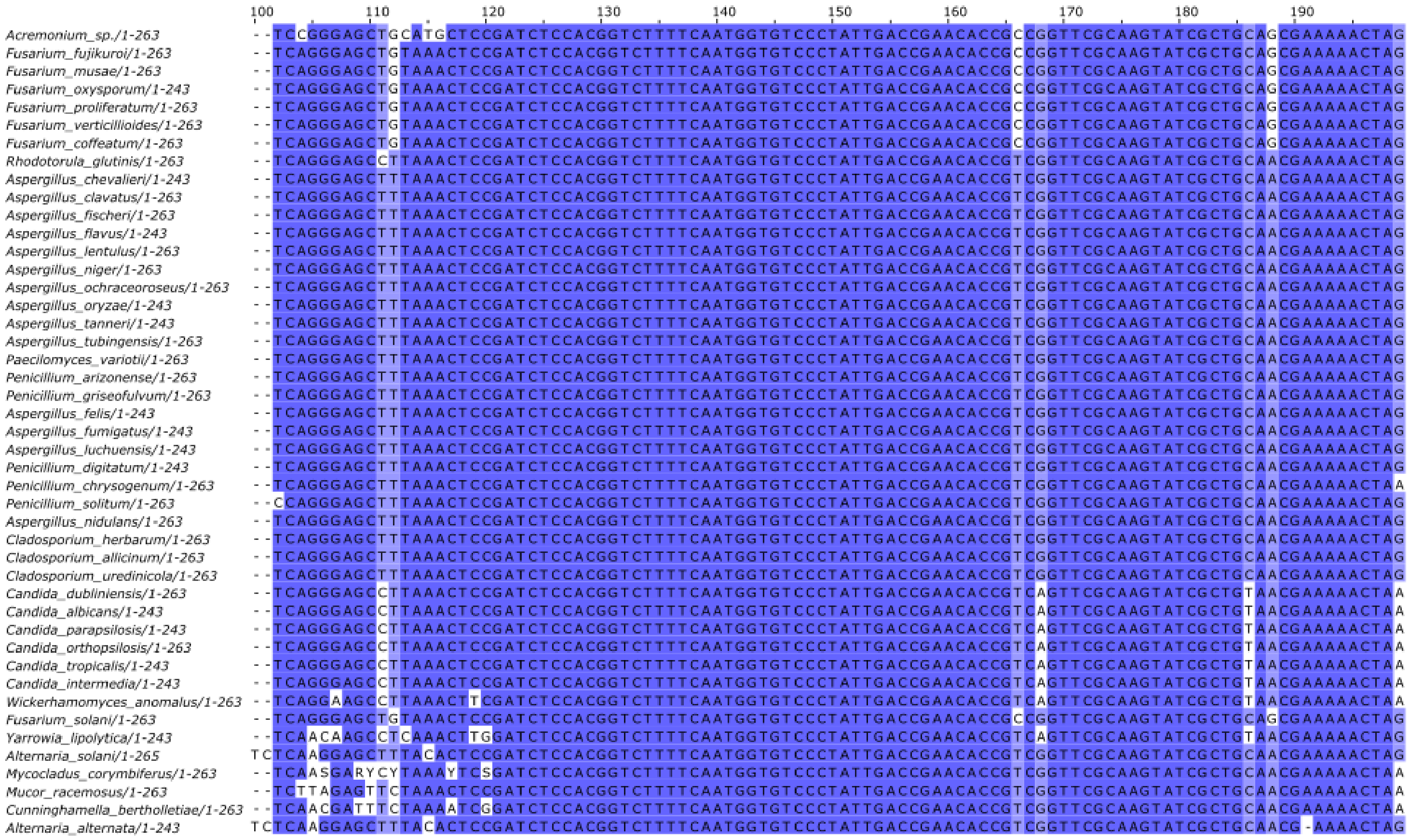
100
110
120
130
140
150
160
170
180
190
Acremonium_sp./1-263
Fusarium_fujikuroi/1-263
Fusarium_musae/1-263
Fusarium_oxysporum/1-243
Fusarium_proliferatum/1-263
Fusarium_verticillioides/1-263
Fusarium_coffeatum/1-263
Rhodotorula_glutinis/1-263
Aspergillus_chevalieri/1-243
Aspergillus_clavatus/1-263
Aspergillus_fischeri/1-263
Aspergillus_flavus/1-243
Aspergillus_lentulus/1-263
Aspergillus_niger/1-263
Aspergillus_ochraceoroseus/1-263
Aspergillus_oryzae/1-243
Aspergillus_tanneri/1-243
Aspergillus_tubingensis/1-263
Paecilomyces_variotii/1-263
Penicillium_arizonense/1-263
Penicillium_griseofulvum/1-263
Aspergillus_felis/1-243
Aspergillus_fumigatus/1-243
Aspergillus_luchuensis/1-243
Penicillium_digitatum/1-243
Penicillium_chrysogenum/1-263
Penicillium_solitum/1-263
Aspergillus_nidulans/1-263
Cladosporium_herbarum/1-263
Cladosporium_allicinum/1-263
Cladosporium_uredinicola/1-263
Candida_dubliniensis/1-263
Candida_albicans/1-243
Candida_parapsilosis/1-243
Candida_orthopsilosis/1-263
Candida_tropicalis/1-243
Candida_intermedia/1-243
Wickerhamomyces_anomalus/1-263
Fusarium_solani/1-263
Yarrowia_lipolytica/1-243
Alternaria_solani/1-265
Mycocladus_corymbiferus/1-263
Mucor_racemosus/1-263
Cunninghamella_bertholletiae/1-263
Alternaria_alternata/1-243

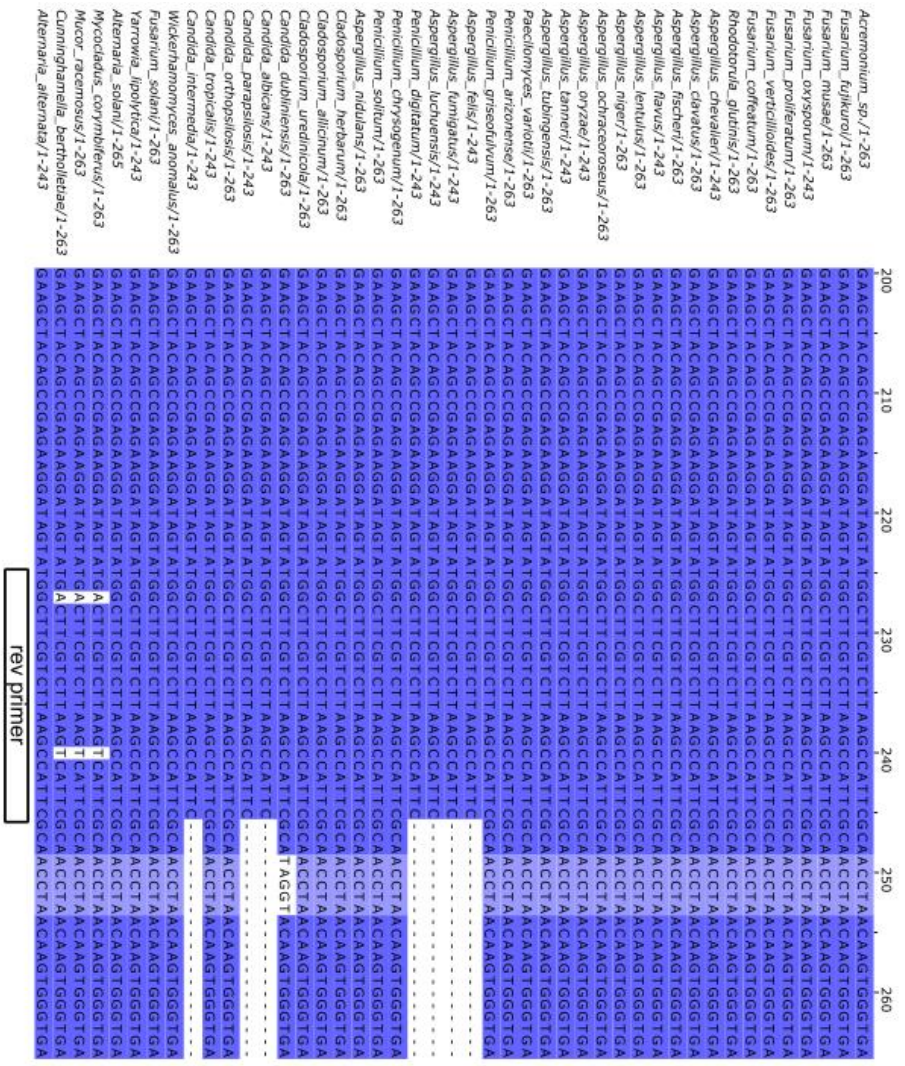


Supplementary Figure 4: Alignment of the fungal rRNA primers and the desired amplicon sequences of tested fungal species. Blue color intensity indicates percentage identity of the sequences. The multiple sequence alignment was generated using MAFFT version 7 and visualized with Jalview 2.11.2.6. All fungal 28S sequences were obtained from GenBank with the indicated reference numbers shown in Supplementary Table 1.

Supplementary Table 1: **Species and sequence information.** The NCBI taxonomy identifiers used for phylogenetic tree generation (Figure 3c) are shown for each species. GenBank reference numbers are displayed for all sequences used to generate the multiple sequence alignments (Supplementary Figure 4). Additionally, information is provided if the primers were reported to be specific for the respective species (by Khot et al.[1]) or if we confirmed primer specificity by PCR (Figure 2b, Figure 3a and b).

| Species | NCBI taxonomy identifier | GenBank reference number | Primer specificity reported in Khot et al.[23] | PCR confirmation |
|---|---|---|---|---|
| *Aspergillus candidus* | 41067 | - | yes | - |
| *Aspergillus chevalieri* | 182096 | NC_057368.1 | - | - |
| *Aspergillus clavatus* | 5057 | NW_001510356.1 | - | - |
| *Aspergillus felis* | 1287682 | CP066505.1 | - | - |
| *Aspergillus fischeri* | 36630 | NW_001510334.1 | - | - |
| *Aspergillus flavus* | 5059 | NC_054697.1 | yes | - |
| *Aspergillus fumigatus* | 746128 | NC_007197.1 | yes | yes |
| *Aspergillus lentulus* | 293939 | NW_022984065.1 | - | - |
| *Aspergillus luchuensis* | 1069201 | NC_054852.1 | - | - |
| *Aspergillus nidulans* | 162425 | KY074658.1 | - | - |
| *Aspergillus niger* | 5061 | MW350015.1 | - | - |
| *Aspergillus oryzae* | 5062 | CP031440.1 | yes | - |
| *Aspergillus tanneri* | 1220188 | NW_022984632.1 | - | - |
| *Aspergillus terreus* | 33178 | - | yes | - |
| *Aspergillus tubingensis* | 5068 | XR_004775245.1 | - | - |
| *Aspergillus ustus* | 40382 | - | yes | - |
| *Candida albicans* | 5476 | CP032012.1 | yes | yes |
| *Candida dubliniensis* | 42374 | AF405231.2 | yes | yes |
| *Candida glabrata* | 5478 | - | yes | yes |

| | | | | |
|---|---|---|---|---|
| *Candida guilliermondii (Meyerozyma guilliermondii)* | 4929 | - | yes | yes |
| *Candida intermedia* | 45354 | LT635764.1 | - | - |
| *Candida kefyr (Kluyveromyces marxianus)* | 4911 | - | yes | yes |
| *Candida krusei* | 4909 | - | yes | - |
| *Candida lipolytica (Yarrowia lipolytica)* | 4952 | CP061014.1 | - | - |
| *Candida lusitaniae* | 36911 | - | yes | - |
| *Candida orthopsilosis* | 273371 | FN812686.1 | - | - |
| *Candida parapsilosis* | 5480 | HE605209.1 | - | - |
| *Candida pelliculosa (Wickerhamomyces anomalus)* | 4927 | NW_017567119.1 | - | - |
| *Candida tropicalis* | 5482 | CP047875.1 | yes | yes |
| *Fusarium coffeatum* | 231269 | NW_022157841.1 | - | - |
| *Fusarium falciforme* | 195108 | - | - | yes |
| *Fusarium fujikuroi* | 5127 | XR_002792831.1 | - | - |
| *Fusarium keratoplasticum* | 1328300 | - | - | yes |
| *Fusarium musae* | 1042133 | NW_025408615.1 | - | - |
| *Fusarium oxysporum* | 5507 | CP052041.1 | - | - |
| *Fusarium petroliphilum* | 203961 | - | - | yes |
| *Fusarium proliferatum* | 948311 | LT841264.1 | - | - |
| *Fusarium solani* | 169388 | FJ345352.1 | yes | yes |
| *Fusarium verticillioides* | 117187 | NW_017387867.1 | - | - |
| *Absidia corymbifera* | 42458 | - | yes | - |
| *Acremonium sp.* | 159075 | GQ867783.1 | - | - |

| | | | | |
|---|---|---|---|---|
| *Penicillium chrysogenum* | 5076 | FJ345355.1 | yes | - |
| *Penicillium griseofulvum* | 5078 | NW_024467492.1 | - | - |
| *Penicillium digitatum* | 36651 | CP060778.1 | - | - |
| *Penicillium solitum* | 60172 | JN642222.1 | - | - |
| *Alternaria alternata* | 5599 | CP061881.1 | - | - |
| *Cunninghamella bertholletiae* | 90251 | FJ345351.1 | yes | - |
| *Mucor racemosus* | 4841 | FJ345353.1 | yes | - |
| *Paecilomyces variotii* | 264951 | FJ345354.1 | yes | - |
| *Rhodotorula glutinis* | 5535 | FJ345357.1 | yes | - |
| *Cladosporium allicinum* | 1338630 | GU214408.1 | - | - |
| *Cladosporium herbarum* | 29918 | GU214410.1 | - | - |
| *Cladosporium uredinicola* | 237183 | EU019264.2 | - | - |
